# svaRetro and svaNUMT: Modular packages for annotation of retrotransposed transcripts and nuclear integration of mitochondrial DNA in genome sequencing data

**DOI:** 10.1101/2021.08.18.456578

**Authors:** Ruining Dong, Daniel Cameron, Justin Bedo, Anthony T Papenfuss

## Abstract

**Background:** The biological significance of structural variation is now more widely recognized. However, due to the lack of available tools for downstream analysis, including processing and annotating, interpretation of structural variant calls remains a challenge.

**Findings:** Here we present *svaRetro* and *svaNUMT*, R packages that provide functions for annotating novel genomic events such as non-reference retro-copied transcripts and nuclear integration of mitochondrial DNA. We evaluate the performance of these packages to detect events using simulations and public benchmarking datasets, and annotate processed transcripts in a public structural variant database.

**Conclusions:** *svaRetro* and *svaNUMT* provide efficient, modular tools for downstream identification and annotation of structural variant calls.

## Findings

### Background

Structural variants (SVs) are polymorphisms and mutations commonly observed in the genome. They range from simple insertions and deletions to complex chromosomal-scale rearrangements. SVs are a significant source of genomic variability in humans, and SV analysis has rapidly become a part of standard pipelines in genomic studies [1–4]. To call SVs from short-read DNA sequencing data derived from individual samples (e.g., germline or cell lines), matched tumour-normal pairs, or multiple related samples, a variety of tools have been developed [5–7].

Interpretation of SV calls requires additional downstream analyses. For example, in tumour genome analysis, annotation of genes disrupted by SVs is a relatively straightforward downstream analysis; due to the splicing logic required, fusion gene prediction is more complicated (e.g. LINX [8]); while inference of chromothripsis [9] or chromoplexy [10]. There are relatively few modular tools available for downstream annotation of SVs. Further complicating this issue, some SV annotation software is highly dependent on specific SV callers (e.g. LINX with GRIDSS) or completely integrated (e.g. AmpliconArchitect [11] and RetroSeq [12]). To cope with the ever-growing number of SV datasets downstream of calling SVs, users need high-quality tools to annotate and understand these calls. Two biological phenomena that are currently underserved by available tools are Nuclear Mitochondrial insertion (NUMTs) [13] and retroposed transcript (RT) insertion.

Nuclear mitochondrial integrations (NUMTs) are formed during mitosis when the nuclear membrane breaks down, allowing mitochondrial DNA (mtDNA) to escape from degrading mitochondria, which is accelerated in cancer, and migrate into the nuclear genome [14,15]. NUMTs are present in the normal genome, having integrated during evolution. Somatic NUMT events have been observed in human cancer cells, but have not been extensively studied, and further investigation is needed to understand their extent and role in cancer development [16,17]. Despite their potential biological significance, these events are often overlooked [18].

Retroposed transcripts (RTs) are associated with LINE element reactivation in cancer [19] but also occur in the germline leading to processed pseudogenes [20]. RTs can interfere with the expression of their parent genes [21], generate antisense transcripts [22], and compete for microRNA binding with their parent genes [23]. Additionally, mutations introduced by the process may drive cancer evolution, particularly when the retroposed transcript is inserted into another gene.

Here, we present two R packages for the downstream analysis of SVs. svaNUMT and svaRetro provide flexible frameworks to analyze and explore NUMTs and RTs. In typical use, SV calls in a VCF file are loaded into a breakend-centric GRanges object using *VariantAnnotation* [24] and *StructuralVariantAnnotation* [25]. The packages then search for evidence supporting events of interest. For RT detection, svaRetro also requires a TxDb object which stores transcript metadata. The TxDb object can be loaded via pre-existing annotation packages or generated from existing data [26]. The output of the packages are lists of GRanges objects that can be converted to various data formats, including BEDPE, supporting further analysis (Figure 1).

**Figure 1.**
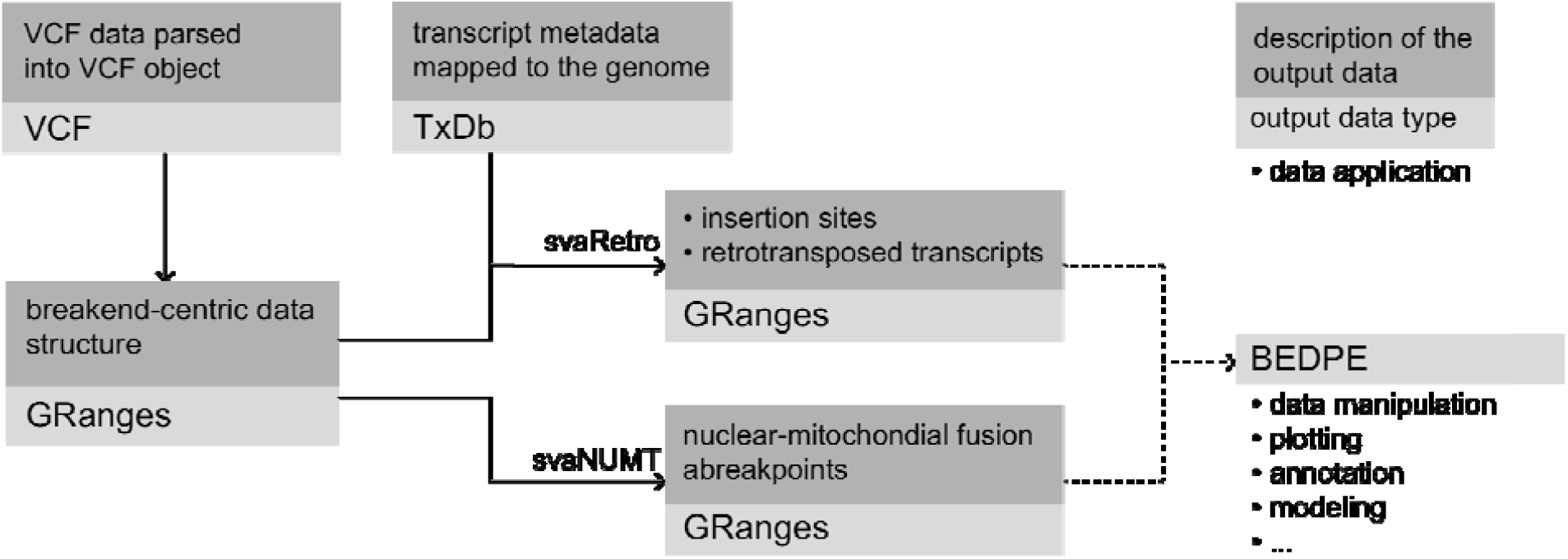
Workflow of svaRetro and svaNUMT. SV calls are first loaded as VCF objects with VariantAnnotation [24], then converted into breakend-centric GRanges with StructuralVariantAnnotation [25]. svaRetro takes as input the Granges data and a TxDb annotation object, which stores the transcript metadata. The output of svaRetro is a list of GRanges grouped by the source gene of the retrotransposed transcripts. svaNUMT requires only the GRanges object as input. The results are grouped by events and the locations of the breakends. The output can be easily converted to BEDPE format, which is commonly used for downstream analyses.

### Input and Output Data Format

Both *svaRetro* and *svaNUMT take as input* a GRanges object with a breakend-centric notation, where a GRanges record is used to represent each breakend, and a breakpoint consists of a pair of breakends. Although breakpoint-centric data structures are available for SV representation (e.g., Pairs object in *rtracklayer* [27]), we have chosen the breakend-centric notation as it simplifies frequent operations in the analysis such as overlap finding with genes and repeats.

The output formats of *svaRetro* and *svaNUMT* are GRanges to support flexible downstream analyses. In cases where alternative formats are required, *StructuralVariantAnnotation* [25] provides functions for format conversion between BEDPE and Pairs [28] with GRanges.

### Identifying Retrotransposed Transcripts

*SvaRetro* identifies RTs using the provided SV calls. RTs are processed transcripts integrated into the genome and characterized by intronic losses and polyadenylation. The candidate insertion sites are scattered across the genome due to the mobilization of transposable elements and are frequently combined with target site duplications (TSD). Therefore, except when the transcript comprises only a single exon, an RT should show a signature of intronic deletions—breakpoints aligned with adjacent exon boundaries from the same mRNA transcript. Additionally, the insertion site is detectable as a rearrangement connecting an exonic edge and a second genomic location (see Figure 2).

**Figure 2.**
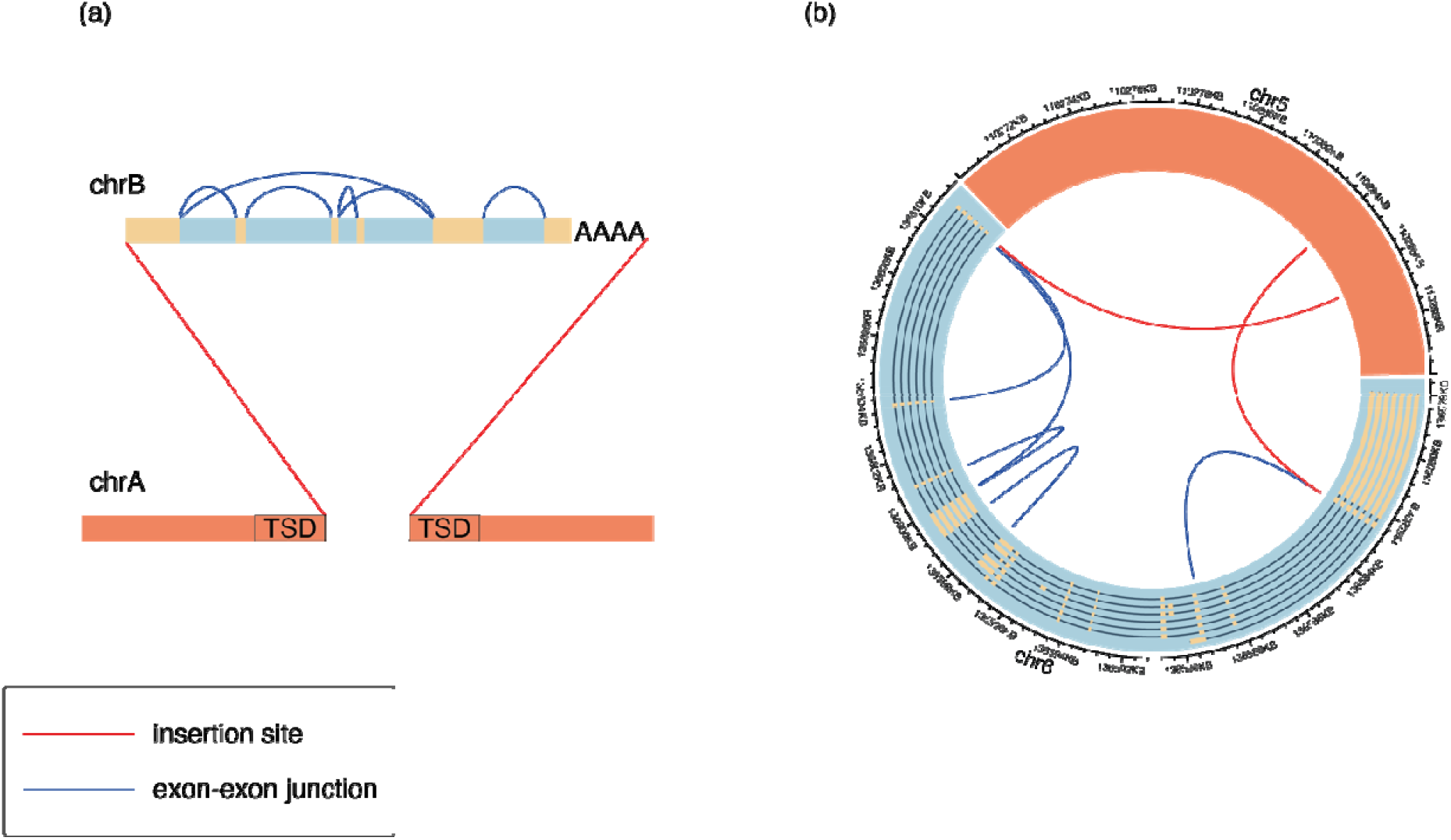
Breakpoint signatures of RT events. (a) A multi-exon RT consists of two breakpoint signatures: exon-exon junctions (blue) and fusions of exon and insertion sites (red). Polyadenylation and target site duplication (TSD) may also be present. (b) A germline BCLAF1 RT on chr5 detected in sample COLO829 [29] with breakpoint calls of exon-exon junctions (blue) and insertion sites (red). Not all exon-exon junctions were detected by the SV caller, and a transitive call connecting exon 1 and exon 4 is also evident.

To detect intronic deletions, overlaps between breakend positions and exon-intron (and intron-exon) boundaries are returned under a maximum gap threshold (default of 100 bps). An RT is reported with higher confidence when more exons are present. This quality, denoted by *minscore*, is evaluated using the proportion of intronic deletions detected from the total possible in the transcript. Depending on the resolution of the SV caller, small exons (e.g. shorter than the read length) can be missed or captured in transitive breakpoints (i.e. a pair of adjacent rearranged segments A-to-B and B-to-C are detected as A-to-C). Meanwhile, breakpoints consisting of an exon boundary and a second genomic location are potential insertion sites. Due to the frequent 5’ truncation in retrotransposons, the maximum gap threshold does not apply here.

The output is a list of GRanges objects consisting of breakend-centric SV calls grouped by the source gene of the retroposed transcript. Each grouped event contains candidate insertion sites and exon-exon junctions, if available. Each candidate insertion site is annotated by the potential source transcript(s) and whether exon-exon junctions are detected for the source transcript(s). Exon-exon junction calls are annotated by the exon indices, corresponding transcripts satisfying the *minscore* threshold, and NCBI gene symbols.

RT insertion sites can be discovered on both 5’ and 3’ sides, only one side, or none. An insertion site could be missed, even when the breakend is reported in the SV callset, due to a sizable 5’ truncation despite the tolerant threshold, a 5’ inversion, or a combination of rearrangements.

### Identifying Nuclear-mitochondrial Genome Fusion Events

*svaNUMT* searches for NUMT events by identifying SVs (in breakend notation) supporting the fusion of nuclear chromosome and mitochondrial genome. In the event of mtDNA integration in nuclear genomes, it is expected that split reads and discordant reads are detected near the integration sites. These features, when picked up by a structural variant caller, are represented as translocation events between mtDNA and nuclear DNA in the SV calls, given the mitochondrial reference genome is included in the library (see Figure 3).

**Figure 3.**
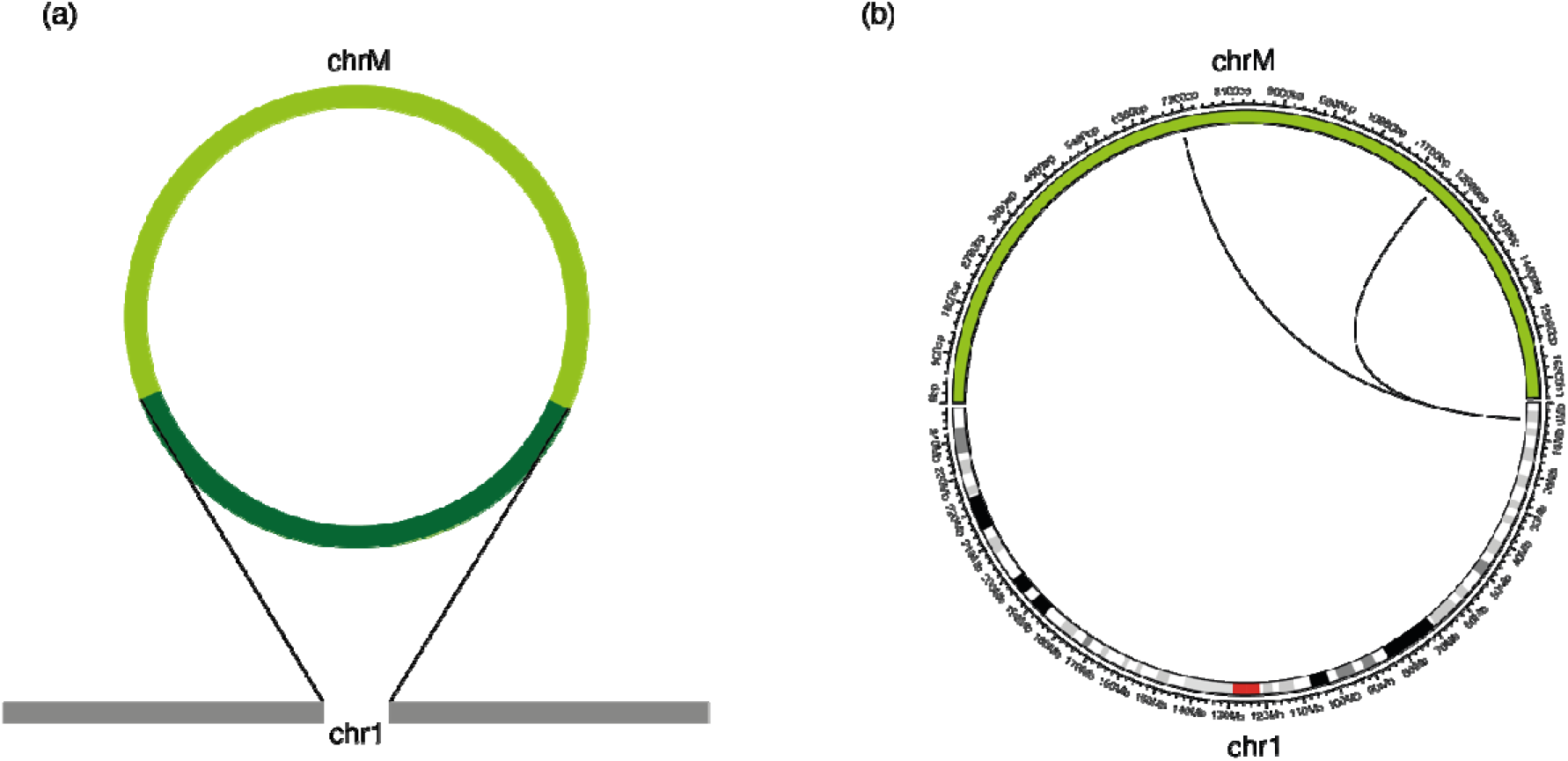
Breakpoint signatures of NUMTs. (a) Schematic of a NUMT event where a sequence from chrM (dark green) is inserted into a nuclear chromosome (chrA). The sequencing read features consists of one breakpoint connecting chrA and chrM for each insertion site. (b) A NUMT event detected in chr1 of the NUMT simulation (described in the text).

A NUMT event consists of two insertion sites, which can be linked by phasing nearby events. *svaNUMT* annotates linked insertion sites where possible. Candidate linked nuclear insertion sites are reported by events as a list of GRanges.

## Benchmarking and Application

### Testing on simulated data

We next tested *svaRetro* and *svaNUMT* using 500 non-overlapping simulated events on chromosome 1. To generate these, chromosome 1 was first divided into 570 uniform intervals. Of these, 507 overlapped (at least 80% overlap) the set of high-confidence Tier 1 regions defined by Zook et al [30]. Intervals not in high confidence regions were excluded. A final set of 500 intervals was then randomly selected. A different random transcript sequence, accompanied by a polyadenylation sequence, was inserted into a random location of each interval. Simulated NUMTs were generated through insertions of 500 mtDNA sequences with polyadenylation on the chr1 sequence, where insertion sites were selected using the same method as described above. The mtDNA sequences included 50 each of lengths 10, 20, 50, 100, 200, 500, 1,000, 2,000, 5,000, and 10,000 base pairs (bp). Paired-end reads at 30x mean coverage were simulated using Art [31] using the HiSeq 2500 error profile for both simulated RTs and simulated NUMTs. SV breakpoints of both samples were called by Manta [6] and GRIDSS [5]. We then used svaRetro and svaNUMT to detect simulated events and analyzed the results with manual inspection.

svaRetro detected 470 out of 500 (94%) RTs from the GRIDSS calls and 443 out of 500 (86%) from Manta calls (candidateSV callset). 23 of the 30 undetected events did not have a breakpoint called by GRIDSS within 100bp of the insertion sites. Out of the seven events where breakends were detected near the insertion sites, 5 events were not mapped to the reference genome, 1 event had secondary calls mapped to an alternative locus, and 1 event was mapped to alternate assembly. Of the 57 undetected events from the Manta calls, 45 did not have a breakpoint called within 100bp of the insertion sites, 11 insertion sequences were not mapped to the reference, and 1 inserted transcript was mapped to an alternate assembly.

svaNUMT detected 276 NUMTs from GRIDSS SV calls and 202 from Manta SV calls. Among the undetected NUMT events, some were due to undetected breakpoints near the insertion sites by the SV callers (14 from GRIDSS and 75 from Manta); some were due to unmapped insertion sequences (122 from GRIDSS and 105 from Manta), where the majority were 10 and 20bp events. In the remainder of the events, only one insertion site breakpoint detected reported (88 from GRIDSS and 118 from Manta). The summary of the results is shown in Figure 4.

**Figure 4.**
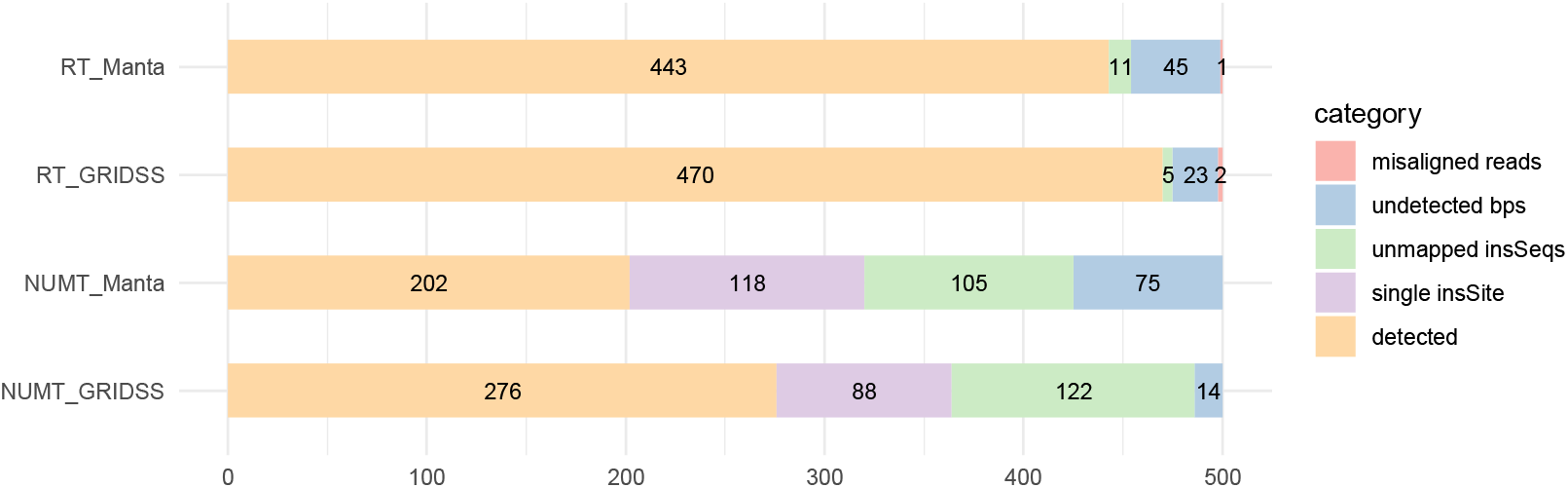
Detection results of the 500 simulated RT and NUMT events by svaRetro and svaNUMT using SV callsets produced by Manta and GRIDSS. For simulated RTs, the undetected events fell into one of categories of unmapped insertion sequences (insSeqs), undetected breakpoints (bps), or misaligned reads. For simulated NUMTs, the unreported events either had one insertion site breakpoint undetected (single insSite), unmapped insSeqs, or undetected bps.

### Benchmarking against existing tool

While there were tools developed for transposable elements from WGS data, the detection of RTs has not been addressed. To our best knowledge, *GRIPper* is the only publicly available RT detection tool [32]. We applied *svaRetro* and *GRIPper* on human germline and tumour datasets, namely HG002, a GIAB cell line using 60× coverage WGS [33], and COLO829, a tumour cell line derived from a cutaneous melanoma [29] with matched lymphoblastoid (normal) cell line. For *svaRetro*, we used GRIDSS to call SVs on these samples. The results were compared with manual inspection. To our knowledge, no publicly available tools exist to detect NUMTs. Consequently, we were unable to benchmark *svaNUMT*.

#### HG002

*GRIPper* reports four instances of RTs in HG002, all of which are detected by *svaRetro* under the same matching threshold of *GRIPper*. In addition, *svaRetro* reports the exon–exon junctions and trace the source of the insertion sequence to the specific transcripts, which is absent in *GRIPper* (Supplementary File 1).

#### COLO829

Three instances of RTs were detected by *GRIPper* in both tumour and matched normal samples. 1 instance of RT is detected in normal sample only (Supplementary File 2 and 3). For one event detected by GRIPper in both the tumour and the normal samples, the inserted transcript sequence mapped to a processed pseudogene (on chr1) as well as the source gene (on chr16). GRIDSS reported this event as a translocation of the in-reference pseudogene from chr1, therefore this event was not reported by svaRetro (see Supplementary Figure 1). The rest RT events were successfully identified by *svaRetro*.

The discrepancy is the result of breakpoint evidence required for RT discovery. *GRIPper* reports putative insertion sites using only split reads, while *svaRetro* collects information on both exon – exon junctions and candidate RT insertion sites, based on breakpoints detected by the structural variant caller of choice.

### Application to gnomAD-SV database

We established a catalogue of non-reference RTs using *svaRetro* on the gnomad-SV dataset [34], where RTs were largely unannotated. NUMT annotation was not applicable as mitochondrial SVs were excluded from the database. In total, 53,529 candidate insertion sites were detected by svaRetro, including events single-exon transcript insertions and/or with insertions with only one side of the insertion detected. The distribution of all source genes and candidate insertion sites are shown in Figure 5. 1298 high-confidence insertion sites were supported by exon-exon junctions and high-quality SV breakpoint calls (using the “PASS” filter). RT insertions can be detected in non-repetitive sequence and across different types of repetitive sequence (Figure 6).

**Figure 54.**
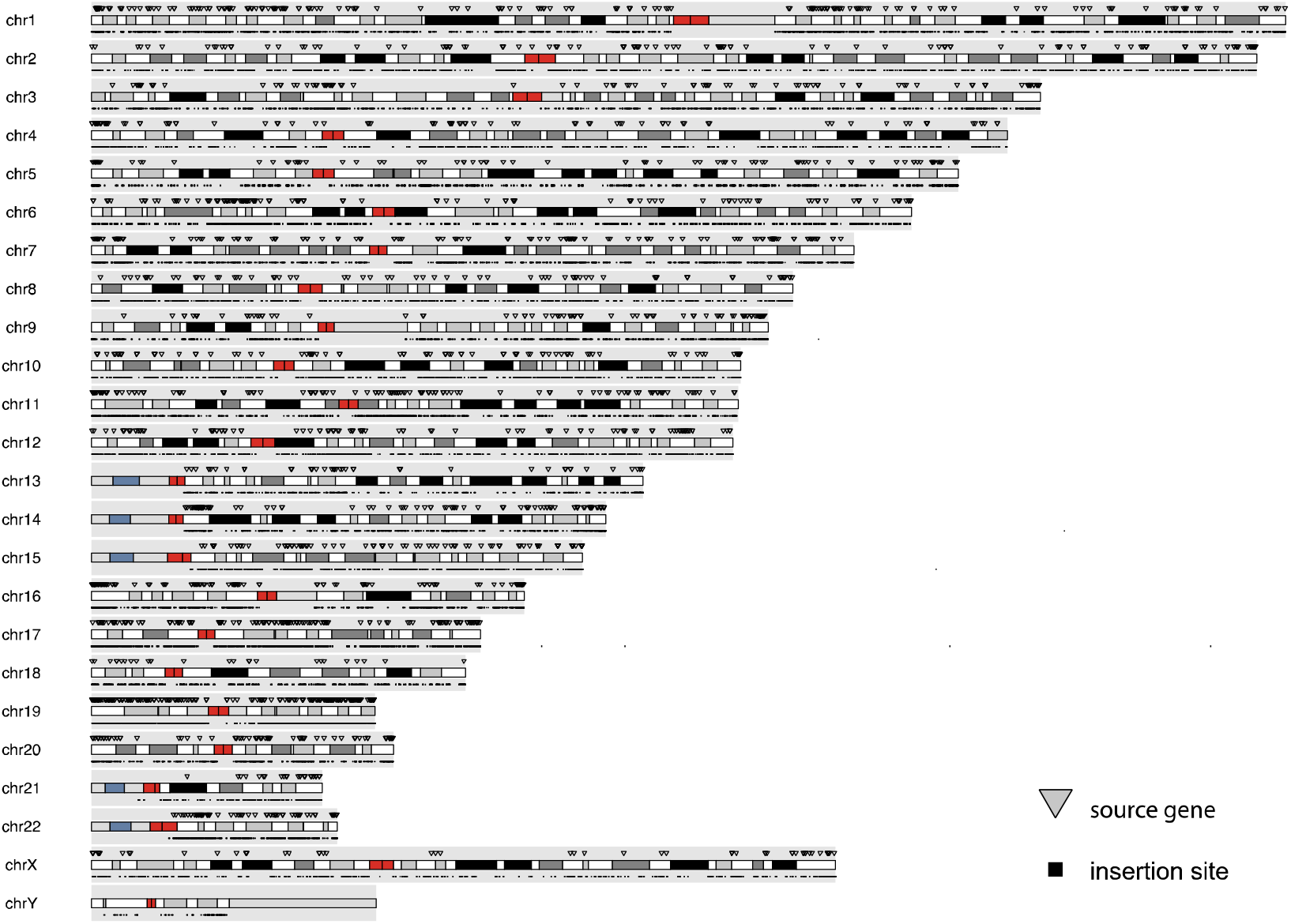
Genomic location of RTs (source gene and insertion locus) detected in gnomAD-SV database.

**Figure 65.**
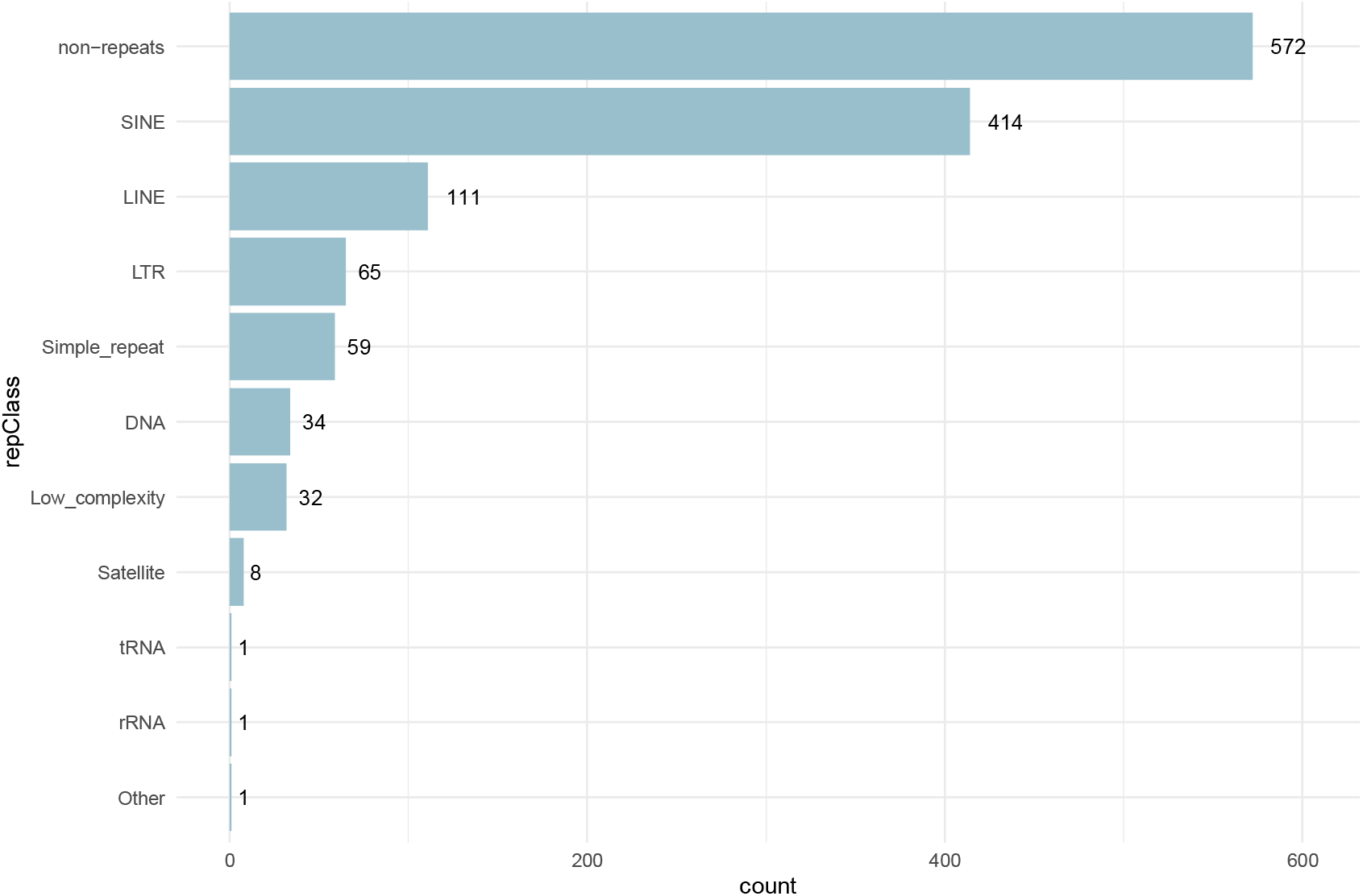
RTs are detectable in gnomAD-SV across different genomic contexts. RepeatMasker [35] annotations of RT insertion regions.

## Conclusion

We present *svaRetro* and *svaNUMT*, R packages developed to facilitate the identification of retro-posed transcript and NUMT insertions. Our tools show outstanding performance on simulation datasets and when benchmarked against existing methods. To further demonstrate its capability, novel RT insertions were discovered by *svaRetro* on a public population SV database. Integrated into the Bioconductor framework, the packages are compatible with many other available tools for more comprehensive downstream analyses.

## Availability of source code and requirements

- Project name: svaRetro and svaNUMT
- Project home page:
  - svaRetro: https://github.com/PapenfussLab/svaRetro
  - svaNUMT: https://github.com/PapenfussLab/svaNUMT
- Operating systems: Platform independent
- Programming language: R
- Other requirements: R ^3^ 4.1, Bioconductor ^3^ 3.14
- License: GPL-3
- RRID:
  - svaRetro: SCR_021380
  - svaNUMT: SCR_021381

## Availability of supporting data and materials

Data supporting the results of this article are available via the GigaScience repository, GigaDB <u>link TBA</u>.

## Declarations

None declared.

## List of abbreviations

SV: structural variant
NUMT: nuclear mitochondrial integration
RT: retroposed transcript
TSD: target site duplication
mtDNA: mitochondrial DNA

## Ethical approval

Not applicable.

## Consent for publication

Not applicable.

## Author contributions

A.T.P. conceived the study. R.D. developed the software, wrote the initial draft of the manuscript. D.C. contributed to the software design. J.B. and A.T.P. oversaw the project. All authors reviewed, contributed to, and approved the manuscript.

