## Supplementary figures and images for "svaRetro and svaNUMT: Modular packages for annotation of retrotransposed transcripts and nuclear integration of mitochondrial DNA in genome sequencing data"

### Supplementary Figure 1

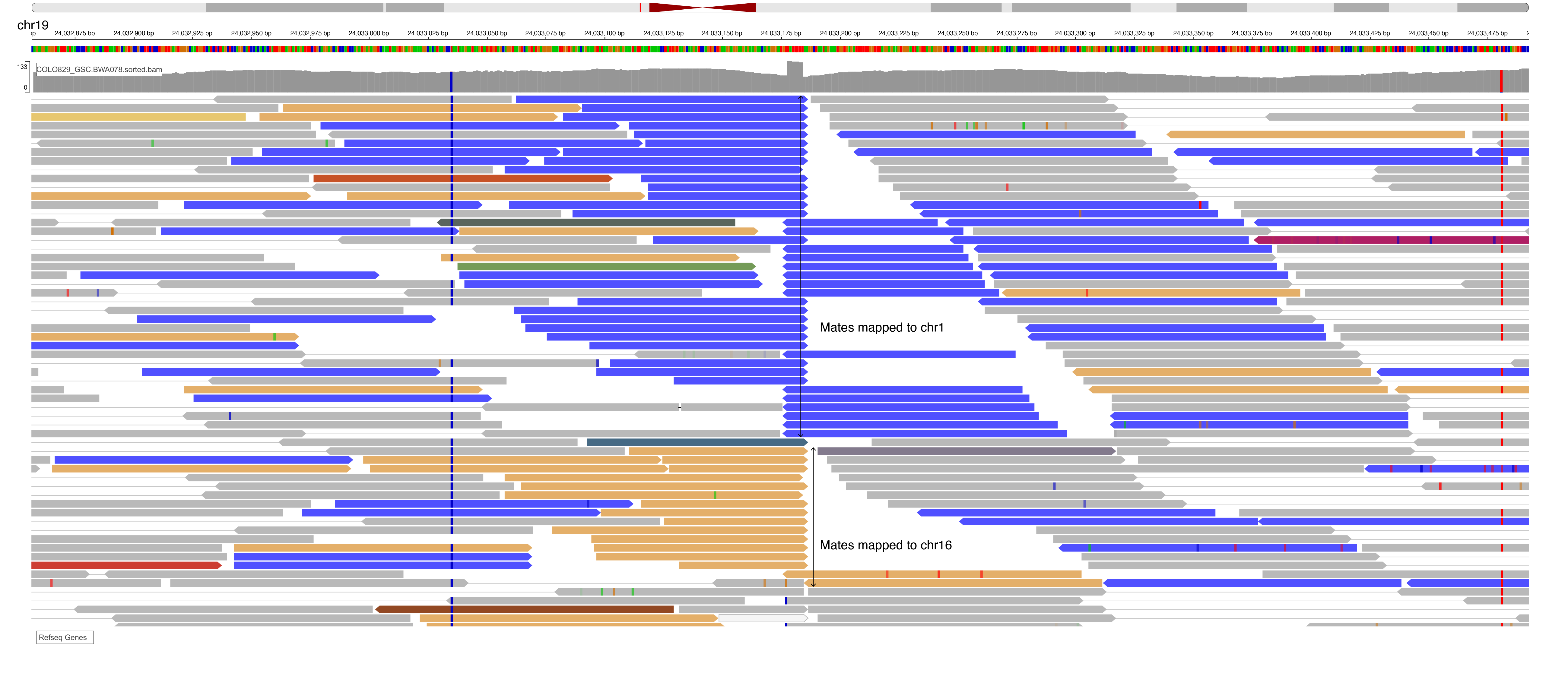
